# A Computational Workflow for Structure-Guided Design of Potent and Selective Kinase Peptide Substrates

**DOI:** 10.1101/2025.07.04.663216

**Authors:** Abeeb A. Yekeen, Cynthia J. Meyer, Melissa McCoy, Bruce Posner, Kenneth D. Westover

## Abstract

Kinases are pivotal cell signaling regulators and prominent drug targets. Short peptide substrates are widely used in kinase activity assays essential for investigating kinase biology and drug discovery. However, designing substrates with high activity and specificity remains challenging. Here, we present Subtimizer (<u>sub</u>strate op<u>timizer</u>), a streamlined computational pipeline for structure-guided kinase peptide substrate design using AlphaFold-Multimer for structure modeling, ProteinMPNN for sequence design, and AlphaFold2-based interface evaluation. Applied to five kinases, four showed substantially improved activity (up to 350%) with designed peptides. Kinetic analyses revealed >2-fold reductions in Michaelis constant (K_m_), indicating improved enzyme-substrate affinity. Two designed peptides exhibited >5-fold improvement in selectivity. This study demonstrates AI-driven structure-guided protein design as an effective approach for developing potent and selective kinase substrates, facilitating assay development for drug discovery and functional investigation of the kinome.

## 1 Introduction

Protein kinases, comprising over 500 members, are central regulators of cellular processes including metabolism, signal transduction, cell growth, and differentiation ^[1–4]^. These enzymes catalyze the transfer of a phosphate group from ATP to specific serine, threonine, or tyrosine residues on their target protein substrates, thereby modulating protein function and complex signaling pathways ^[4–6]^. Consequently, the dysregulation of kinase activity is implicated in numerous diseases, including cancer, inflammatory disorders, and metabolic syndromes, making kinases important targets for both fundamental research and therapeutic intervention ^[2, 7]^. Over 85% of the human kinome is implicated in various diseases, establishing kinases as one of the most important and extensively pursued classes of drug targets ^[4, 8, 9]^.

This has led to the development and clinical success of numerous kinase inhibitors, with over 100 small-molecule inhibitors approved for clinical use ^[6, 8]^. However, these drugs target only about 10% of the human kinome, with the majority belonging to the tyrosine kinase family, leaving most kinases underexplored and underutilized in clinical contexts ^[8–11]^. Robust kinase activity assays are indispensable tools for studying kinase biology and driving drug discovery efforts ^[12, 13]^, yet, more than 50% of the over 500 known kinases do not have established high-throughput assays due to the lack of necessary tools, including optimal and assay-suitable substrates ^[3, 7, 14–16]^. This lack of assays hinders research on these kinases and the development of potential therapeutics targeting them.

In practice, researchers often use short synthetic peptide substrates to quantify kinase activity, enabling the study of kinase function and inhibitor development ^[17–19]^. Compared to full-length protein substrates, peptides offer several advantages, including ease of synthesis, purification, and storage. They are also more cost-effective and readily adaptable to various assay formats, including high-throughput screening ^[7, 17–19]^. However, most existing kinase substrates are either promiscuous or non-selective leading to high background and limited specificity, and hampering accurate activity measurement, especially for complex mixtures like cell lysates or tissue extracts ^[7, 18, 20, 21]^. Traditional substrate discovery and optimization methods, such as positional scanning peptide libraries or phosphoproteomic profiling, are costly, time-consuming and labor-intensive ^[15, 19, 22, 23]^. These methods often involve synthesizing libraries of peptides with systematic substitutions, followed by experimental determination of phosphorylation efficiency. While these approaches have been successful in identifying promiscuous substrates or minimal recognition motifs surrounding phosphosites, they are often limited by their inability to fully explore the vast sequence space beyond simple substitutions ^[15, 23]^. Thus, scalable, generalizable approaches to kinase substrate design and optimization are needed to cater to the growing need for tools to explore uncharacterized kinases, elucidate new mechanisms, and support kinase-targeted drug discovery ^[9]^.

Existing computational efforts have primarily focused on identifying phosphorylation sites on protein substrates ^[24, 25]^, predicting kinase-protein substrate pairs ^[26]^, or predicting kinases responsible for known phosphosites ^[15, 27]^. Despite the long-standing interest and practical importance of designing optimal peptide substrates for kinases ^[28]^, computational design strategies have remained largely unexplored ^[24, 25]^. One notable existing design method is KINATEST-ID, developed to generate peptide substrates for use in chelation-enhanced fluorescence (CHEF) assays for tyrosine kinases ^[17]^. However, KINATEST-ID’s reliance on kinase compatibility with phosphorylation-dependent lanthanide ion coordination (specifically terbium, Tb^3+^) limits its generalizability to diverse assay formats beyond Tb^3+^-sensitization CHEF assays ^[17]^.

Recent advances in AI-based protein modeling methods—propelled by the groundbreaking achievement of AlphaFold2 (AF2) for protein structure prediction ^[29]^ and followed by other related methods including RosettaFold ^[30]^, ESMFold ^[31]^, and AF-Multimer ^[32]^—have revolutionized structural biology and protein engineering ^[33–35]^. Additionally, the field of protein sequence design (predicting amino acid sequences that fold into a given protein structure) has also seen a rapid emergence of innovative AI models with exceptional performance such as ProteinMPNN ^[36]^, ESM-IF1 ^[37]^, ABACUS-R ^[38]^, and more. The transformative impact of these advances was recognized with the 2024 Nobel Prize in Chemistry awarded to David Baker for computational protein design and jointly to Demis Hassabis and John Jumper for protein structure prediction ^[39–41]^.

In this study, we define the design of optimal peptide substrates for kinases as a protein design problem and present subtimizer, a pipeline that integrates Nobel Prize-winning AI-based protein design and structure prediction methods. We previously demonstrated that established protein design tools, such as the ABACUS2 statistical energy function ^[42, 43]^ and the RosettaDesign physics-based method ^[44, 45]^ can reprogram protease-substrate interactions ^[46]^. We hypothesized that recent advances in AI-based protein modeling and design would overcome the limitations of physics-based and statistically-learned empirical energy functions in selectively optimizing enzyme-substrate interactions ^[46]^. To this end, we describe a streamlined structure-guided kinase substrate design workflow that utilizes AF-Multimer to predict the structure of a kinase in complex with a starting peptide substrate, ProteinMPNN for sequence optimization, and AF2-based interface prediction and evaluation ^[47]^. As a proof of concept, we optimized known but suboptimal peptide substrates for a set of kinases, and the resulting designed peptides were experimentally evaluated.

## 2 Results

### 2.1 An AI-driven computational pipeline for designing optimal kinase peptide substrates

The subtimizer pipeline Figure 1A begins with the amino acid sequences of a target kinase and a starting peptide substrate, which could be a known, literature-reported, or kinase family-related substrate sourced from a phosphoproteomic database. Multiple 3D structure models of the kinase-peptide complex are then predicted using AF-Multimer. Given that the accuracy of the predicted complex structure is critical for the success of subsequent design steps, we implemented a filtering step based on the interface Predicted Template Modeling (ipTM) score. The top 5 complexes with ipTM > 0.75, indicating high-confidence prediction of the kinase-peptide interface, were retained for further processing. The predicted complex structures serve as input for the fixed-backbone ProteinMPNN design step, where novel peptide sequences are generated on the backbone of the peptide in each complex. The amino acid identity of the phosphosite (Ser/Thr/Tyr) and kinase sequence are fixed. The goal is to generate novel sequences predicted to optimize the interaction of the complex, with the assumption that improved binding would potentially enhance catalysis.

**Figure 1:**
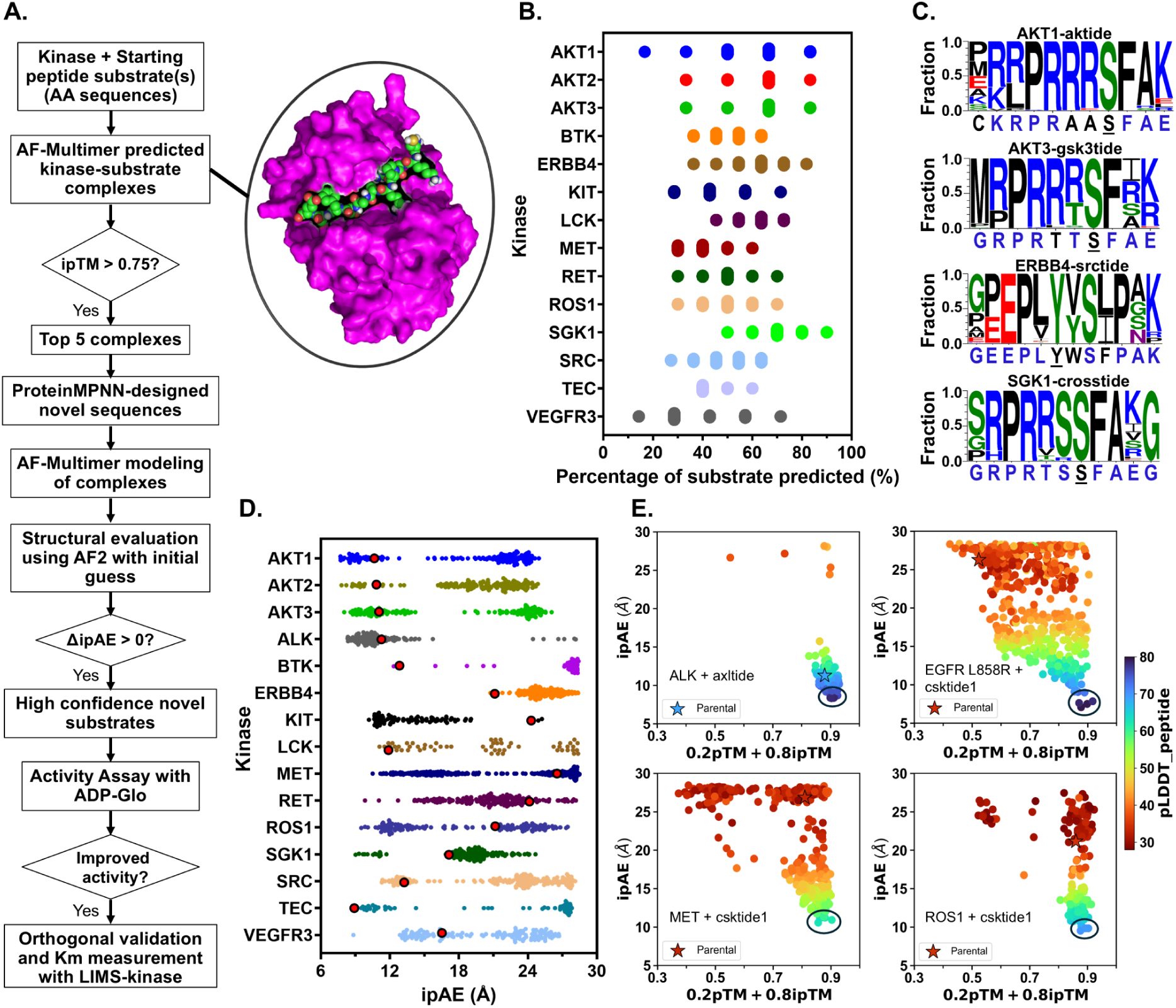
Computational workflow and validation of the Subtimizer pipeline for kinase peptide substrate design. (**A**) Schematic overview of the Subtimizer computational pipeline integrating AlphaFold-Multimer, ProteinMPNN, and AlphaFold2-based structural evaluation for kinase substrate optimization. **B**) Percentage of parental substrate sequence recovered by designed peptides across representative kinase-substrate pairs, demonstrating the pipeline’s ability to capture known functional features. (**C**) Sequence logos showing amino acid preferences at each position for designed peptides compared to parental substrates for selected kinase-substrate pairs. (**D**) Distribution of interface predicted aligned error (ipAE) scores for designed peptides (colored dots) versus parental substrates (bigger red dots) across all tested kinases, with lower scores indicating better predicted binding (interface) quality. (**E**) Correlation plots between ipAE scores and combined pTM and ipTM confidence metrics (0.2 *·* pTM + 0.8 *·* ipTM) for representative kinase-substrate pairs, with peptide confidence (pLDDT) shown by color gradient. In each plot, the starting peptide is shown as a starred dot. Encircled dots are the highest confidence designed peptides from which sequences were selected for experimental testing.

The newly generated peptide sequences are then paired with the kinase sequence and subjected to a second round of kinase-peptide structure modeling using AF-Multimer. The modeled complexes are then evaluated for kinase-peptide interface and binding quality using a modified version of AF2 (AF2 with initial guess) described by Bennett et al. (2023) ^[47]^. Designed peptides with average predicted aligned error of interchain residue pairs (ipAE) score *≤* 10 are considered high-confidence binders ^[47]^. When ΔipAE *>* 0 relative to the parent peptide, they are considered computationally improved. Additional evaluation metrics utilized include peptide pLDDT (predicted local distance difference test) ^[29]^ and a weighted sum of pTM and interface pTM (0.2 *·* pTM + 0.8 *·* ipTM) ^[32]^.

To validate this workflow, we applied it to 47 kinase-substrate pairs experimentally validated in-house. This set comprised 25 unique kinases, each paired with one or more validated but suboptimal peptide substrates. We evaluated how well the Subtimizer workflow recovers residues present in the experimentally validated substrates. Figure 1B shows the percentage substrate recovery for a select set of kinases (see Figure S1 for all pairs). Figure 1C shows sequence logos depicting amino acid preferences for the designed peptides for a few representative kinase-substrate pairs (see Figure S2 for all pairs). Predictions for over half of the evaluated kinase-peptide pairs successfully recovered at least 70% (and up to 90% in some cases) of the residues in the validated substrates (Figures S1 and S2), demonstrating the pipeline’s ability to generate sequences with features consistent with known substrates.

For each kinase-peptide pair, structural evaluation metrics, including ipTM and ipAE scores, were calculated using AF-Multimer and AF2 with-initial-guess ^[47]^. The distribution of ipAE scores for representative kinase complexes of the designed and parental peptides is shown in Figure S1D (see Figure S3 for all kinase-peptide pairs). For all 47 kinase-peptide pairs except SGK1-1aktide, there are designed peptide complexes with ipAE lower than that of the parent peptide (Figure 1D and S3). In addition, most kinases have designed peptides with ipAE < 10 for at least one substrate type. As shown in Figures 1E and S4, lower ipAE scores correlate with high ipTM and pLDDT scores, indicating that these metrics are suitable for evaluating prediction confidence. These structural evaluation metrics were used to filter the designed sequences down to a smaller set of high confidence designs to be prioritized for experimental testing.

### 2.2 Designed Peptide Substrates Showed Improved Potency In Vitro

To experimentally evaluate the computationally optimized peptide sequences, we chose five kinases based on commercial availability. Using the ipAE, pLDDT, and ipTM scores, we ranked the Subtimizer-designed sequences, and for each kinase we selected two to five peptides for synthesis and experimental testing. We evaluated the activity of the peptides using the ADP-Glo kinase assay, a luminescence-based method that quantifies ADP production as a measure of kinase activity. First, we measured the ADP-Glo signals for the parental peptide and the baseline (no enzyme background) for the five kinases. As shown in Figure 2A, we established that all five kinases are active under the assay conditions and that the signal generated from the phosphorylation reaction is significantly above background noise. The varying signal intensities across different kinases reflect differences in their specific activities and the efficiency of the parent peptide substrates.

**Figure 2:**
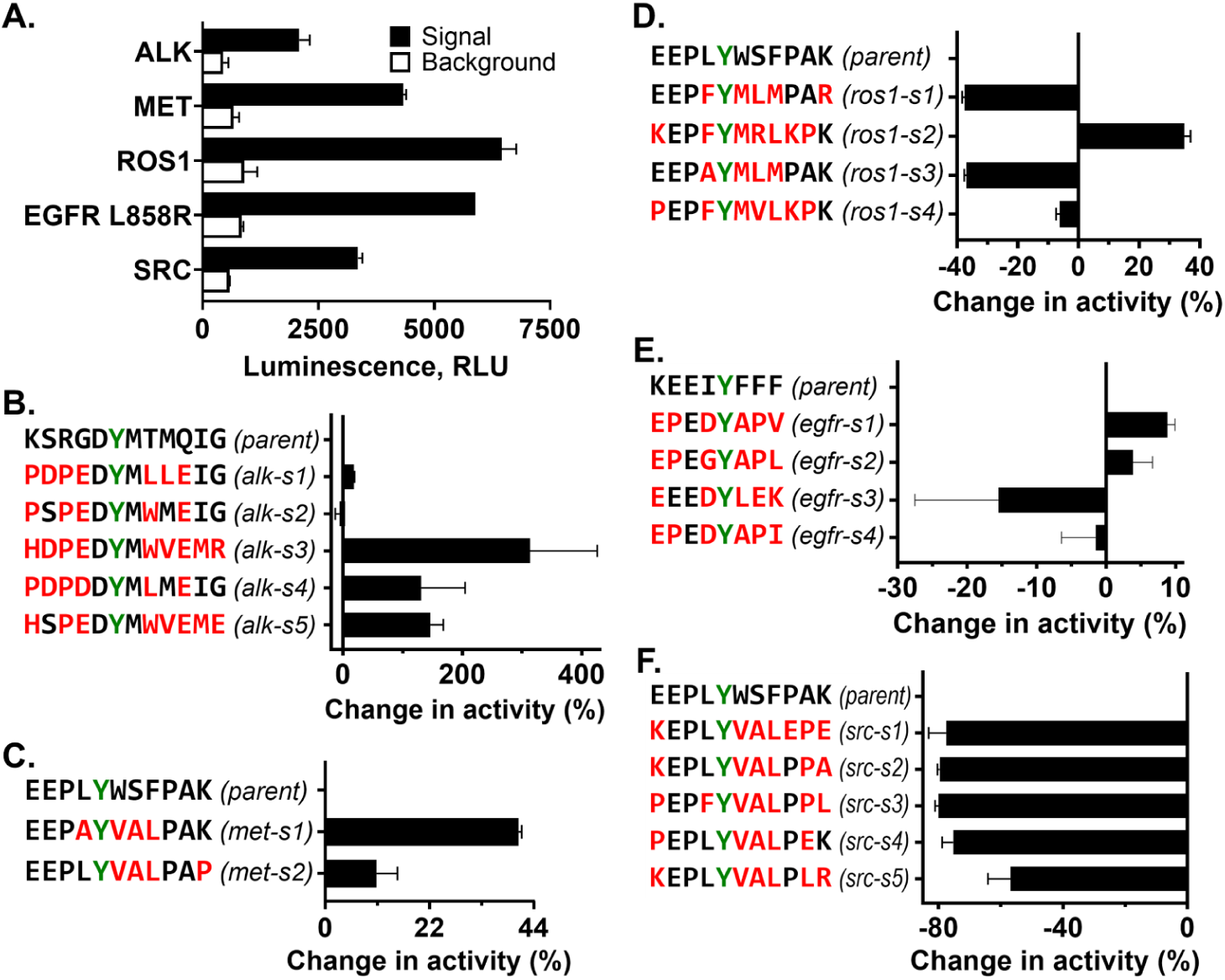
Experimental validation of designed peptides shows improved kinase activity. (**A**) ADP-Glo assay baseline measurements showing kinase activity with parental substrates and background controls for all five tested kinases. (**B–F**) Relative percentage change in kinase activity for designed peptides compared to parental substrates for ALK (**B**), MET (**C**), ROS1 (**D**), EGFR L858R (**E**), and SRC (**F**). Designed peptides are color-coded with sequence information provided. Phosphosite residues that were fixed during design are highlighted in green. Four out of five kinases showed substantial activity improvements with at least one designed peptide.

##Figures 2B-F show, for ALK, MET, ROS1, EGFR, and SRC, respectively, the relative percentage change in kinase activity for the designed peptides compared to their respective parent peptides. As shown in Figure 2B, four out of five peptides designed for ALK showed improved activity compared to the parent peptide KSRGDYMTMQIG. Notably, peptide alk-s2 (PSPEDYMWMEIG) demonstrated a remarkable increase in activity, exceeding 300% relative to the parental axltide substrate. Alk-s5 (HSPEDYMWVEME) and alk-s4 (PDPDDYMLMEIG) also showed substantial improvements of 146% and 130%, respectively. For MET, two designed peptides met-s1 (EEPAYVALPAK) and met-s2 (EEPLYVALPAP) were tested as shown in Figure 2C. Compared to the parent peptide EEPLYWSFPAK, both met-s1 and met-s2 exhibited enhanced activity, showing relative improvements of approximately 41% and 11%, respectively.

For ROS1 (Figure 2D), only ros1-s2 (KEPFYMRLKPK) showed improved activity relative to the parent EEPLYWSFPAK, demonstrating an increase of about 35%. Evaluation of designed substrates for EGFR L858R (Figure 2E) revealed that two out of four peptides, egfr-s1 (EPEDYAPV) and egfr-s2 (EPEGYAPL), showed improved activity compared to the parent peptide KEEIYFFF. Egfr-s1 exhibited an increase in activity of about 10%, while egfr-s2 showed approximately 5% increase. Unlike ALK, MET, ROS1, and EGFR L858R, none of the five designed peptides tested for SRC activity (Figure 2F) showed better activity compared to the parent peptide. Rather, all designed SRC peptides exhibited reduced activity, with decreases ranging from approximately 57% to 80%. Overall, experimental validation using the ADP-Glo assay confirmed that the Subtimizer pipeline can design peptide substrates with substantially improved activity. The tests also showed the high success rate of the pipeline, as optimized peptides can be identified by experimentally testing as few as two peptides.

### 2.3 Improved Activity of Designed Peptides Correlates with Enhanced Binding Affinity

To further characterize the improved designed peptide substrates and gain insights into the mechanism underlying their enhanced activity, we determined the Michaelis constant (K_m_) for the most potent designed peptides and their corresponding parent substrates using the mass spectrometry-based LIMS-kinase assay previously developed in our group ^[16]^. Figures 3A and 3B illustrate the schematic of a kinase assay reaction and the multiple reaction monitoring (MRM) phosphorylation detection system employed in the LIMS-kinase assay. The MRM setup comprises a RapidFire liquid chromatography sampler (Agilent) coupled to a triple quadrupole mass spectrometer (AB Sciex 6500). Quadrupole 1 (Q1) acts as a mass filter, selecting precursor ions based on their mass-to-charge ratios. Q2 functions as a collision cell, fragmenting these precursor ions into product ions. Q3 serves as an additional mass filter, specifically monitoring desired product ion fragments originating from the Q1-selected parent ion (Figure 3B).

**Figure 3:**
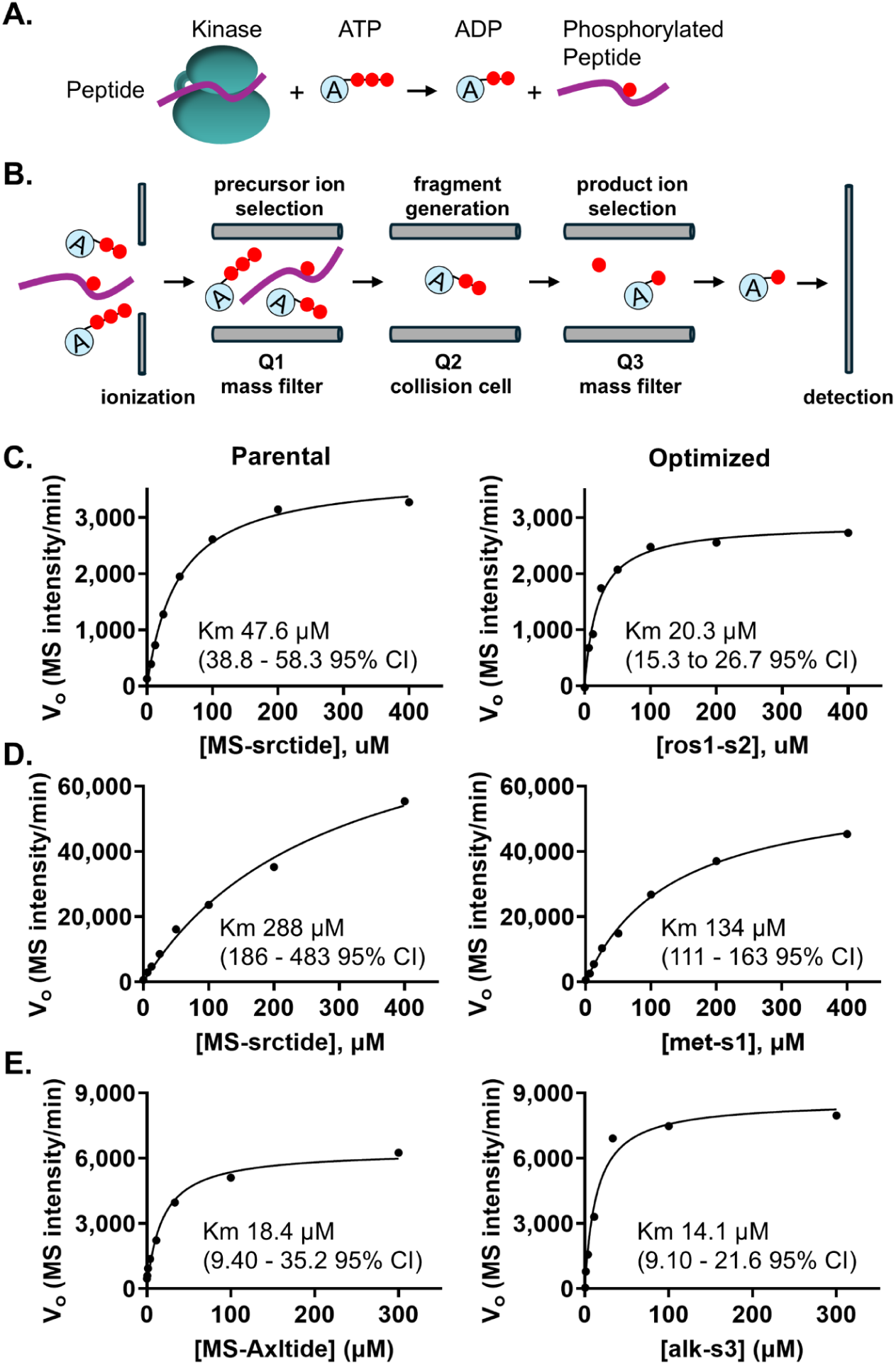
Kinetic characterization reveals improved binding affinity of designed peptides. (**A**) Schematic of kinase phosphorylation reaction monitored by the LIMS-kinase assay. (**B**) Multiple reaction monitoring (MRM) setup showing the mass spectrometry detection pathway for ADP quantification. (**C–E**) Michaelis-Menten kinetic analysis comparing parental and optimized peptides for ROS1 (**C**), MET (**D**), and ALK (**E**). Left panels show time-course data at varying substrate concentrations; right panels show saturation curves with fitted K_m_ values and 95% confidence intervals. The designed peptides demonstrated lower K_m_ values, indicating improved apparent binding affinity.

To determine kinase activity, we directly measured ADP production by following specific ADP ion fragments rather than quantifying phosphorylated peptide fragments as in our original published LIMS-kinase assay ^[16]^. We chose to measure ADP production rather than the phosphorylated peptide product to avoid potential variations in ionization efficiency or fragmentation patterns that can arise from differences in amino acid sequences and physicochemical properties of different peptide substrates. This ensures not only a more direct and unbiased comparison of potency across different peptides, but also broad applicability of optimized substrates across different assay platforms. Initial tests confirmed that we were able to selectively detect an ADP fragment in kinase/peptide reactions (Figure S5). The ADP signal was enzyme concentration- and time-dependent, consistent with the expected response of a functional assay. We then optimized the LIMS-kinase assay for ROS1, MET, ALK, and EGFR L858R. We evaluated two enzyme concentrations with the objective of selecting the concentration where activity was linear to 60 minutes with a strong signal (Figure S5).

We ran LIMS-kinase assays using a range of peptide concentrations. We plotted the initial reaction rate for each concentration to generate Michaelis-Menten curves. The curves were used to calculate K_m_ values for parental and most potent Subtimizer-designed peptides for ROS1, MET, and ALK, respectively (Figures 3C-E, Figures S6-S8). We could not run a successful K_m_ study with EGFR L858R due to the insolubility of the parental MS-csktide1 peptide at high concentrations, similar to our previous observation ^[16]^. For ROS1 (Figure 3C, Figure S6), the parent peptide (MS-srctide) exhibited a K_m_ of 47.6 *µ*M while the optimized peptide ros1-s2 showed a significantly lower K_m_ of 20.3 *µ*M. This more than two-fold reduction in K_m_ indicates that the designed peptide has a substantially higher apparent affinity for ROS1 compared to the parent substrate.

While the parent peptide substrate MS-srctide had a K_m_ of 288 *µ*M for MET (Figure 3D, Figure S7), the designed peptide met-s1 showed an improved activity with a K_m_ of 134 *µ*M—an over 2-fold reduction in K_m_ for MET. For ALK (Figure 3E, Figure S8), the parent peptide MS-axltide exhibited a K_m_ of 18.4 *µ*M while the designed peptide alk-s3 showed a lower K_m_ of 14.1 *µ*M. This decrease in K_m_ suggests improved apparent affinity of the designed peptide for ALK, consistent with the enhanced activity observed in the ADP-Glo assay. The consistent observation of lower K_m_ values for the Subtimizer-designed peptides with improved activity across multiple kinases provides strong evidence that the computational pipeline optimizes substrates for kinase-substrate interaction, generating peptide sequences that bind more tightly to the kinase active site.

### 2.4 Subtimizer-Optimized Substrates Showed Improved Selectivity for Target Kinases

Beyond improving activity, the enhancement of substrate selectivity for a target kinase over other kinases is critically important in practical applications such as assays in complex environments like cells or cell lysates. This is particularly relevant for kinases with similar substrate recognition motifs or those belonging to the same family. To evaluate the selectivity of the Subtimizer-designed peptides, we assessed the activity of selected optimized peptides against both their intended target kinases and a related kinase that utilizes the same parent peptide. The MS-srctide peptide (EEPLYWSFPAK) is a substrate for many kinases including MET and ROS1 ^[16]^, and it served as the parent substrate for both the met-s1 peptide, optimized for MET, and the ros1-s2 peptide, optimized for ROS1 (Figure 4A). This common parentage provides a direct context for evaluating whether the design process introduced kinase-specific selectivity. Figure 4B shows the activity of MET kinase with each of the designed peptides ros1-s2 and met-s1, as measured by the LIMS-kinase assay (quantifying ADP production). The MET kinase exhibited high activity with met-s1 peptide (designed for MET) as expected and consistent with the improved activity observed in the ADP-Glo assay (Figure 2C). In contrast, MET kinase showed over 4-fold reduction in activity with the ros1-s2 peptide (designed for ROS1). This indicates that the modifications introduced during optimization of ros1-s2 for ROS1 reduced its efficiency as a substrate for MET, even though both peptides originated from the same MS-srctide parent.

**Figure 4:**
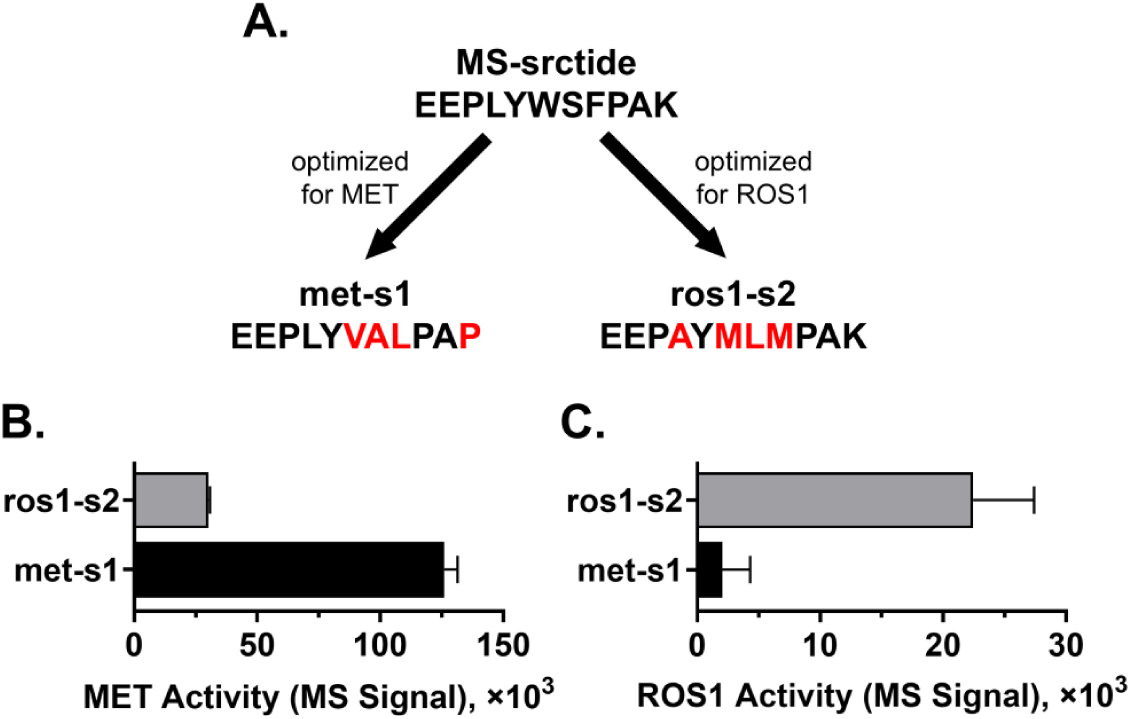
Designed peptides exhibit enhanced selectivity for target kinases. (**A**) Schematic showing the common parental substrate MS-srctide optimized separately for MET (met-s1) and ROS1 (ros1-s2). (**B**) MET kinase activity with designed peptides ros1-s2 and met-s1, showing preferential activity with MET-optimized peptide. (**C**) ROS1 kinase activity with the same peptides, demonstrating preferential activity with ROS1-optimized peptide. The reciprocal selectivity pattern confirms successful kinasespecific optimization.

Conversely, when the same two peptides ros1-s2 and met-s1 were incubated with ROS1 kinase (Figure 4C), ROS1 kinase showed an over 11-fold reduction in activity with the met-s1 peptide (designed for MET) compared to the ros1-s2 peptide. However, with the ros1-s2 peptide, ROS1 demonstrated high activity as anticipated from the ADP-Glo activity data (Figure 2D) and the K_m_ data (Figure 3C). This reciprocal pattern of activity demonstrates that the Subtimizer design process successfully introduced selectivity into the designed peptides. The MET-optimized peptide (met-s1) is preferentially phosphorylated by MET, while the ROS1-optimized peptide (ros1-s2) is preferred by ROS1, despite their common origin from the less selective MS-srctide peptide.

### 2.5 Structural and Computational Analyses of Optimized Kinase-Peptide Complexes

To gain deeper understanding of the molecular mechanisms underlying the observed improvements in activity, binding affinity (K_m_), and selectivity of the designed peptide substrates, we performed structural and computational analyses of the predicted kinase-peptide complexes of the most potent peptides compared to the parental peptides. We performed a physics-based refinement of the AF-Multimer models of the complexes using Rosetta FlexPepDock ^[48]^. The FlexPepDock is a high-resolution, sub-angstrom quality, protein-peptide docking protocol implemented as a module within the Rosetta framework and is capable of refining protein-peptide complex models to near-native structures ^[48]^. We analyzed the kinase-peptide interactions observed in the refined structures (Figure 5). We also analyzed the computed Rosetta energy and interface scores of the refined structures to quantitatively assess the predicted binding interactions (Table 1).

**Figure 5:**
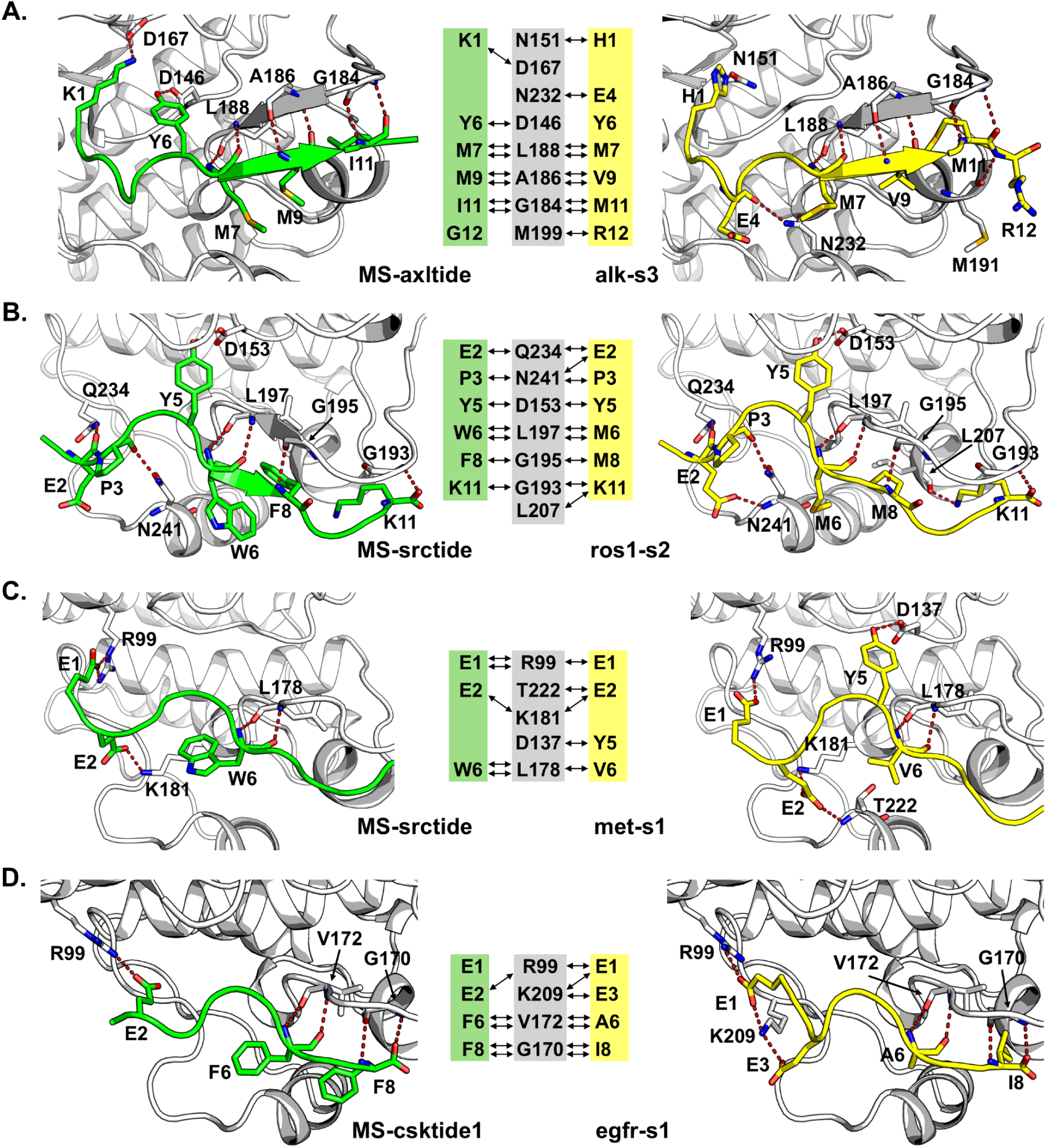
Structural analysis reveals molecular basis for improved activity. (**A-D**) Comparison of AlphaFold-Multimer predicted structures and hydrogen bonding networks for parental (green) versus designed (yellow) peptides in complex with ALK (**A**), ROS1 (**B**), MET (**C**), and EGFR L858R (**D**). Kinase structures are shown in gray ribbon representation. Green boxes highlight hydrogen bond interactions formed by parental peptides; yellow boxes show interactions formed by designed peptides. Designed peptides generally form additional or optimized hydrogen bonds that correlate with improved experimental activity.

**Table 1:**
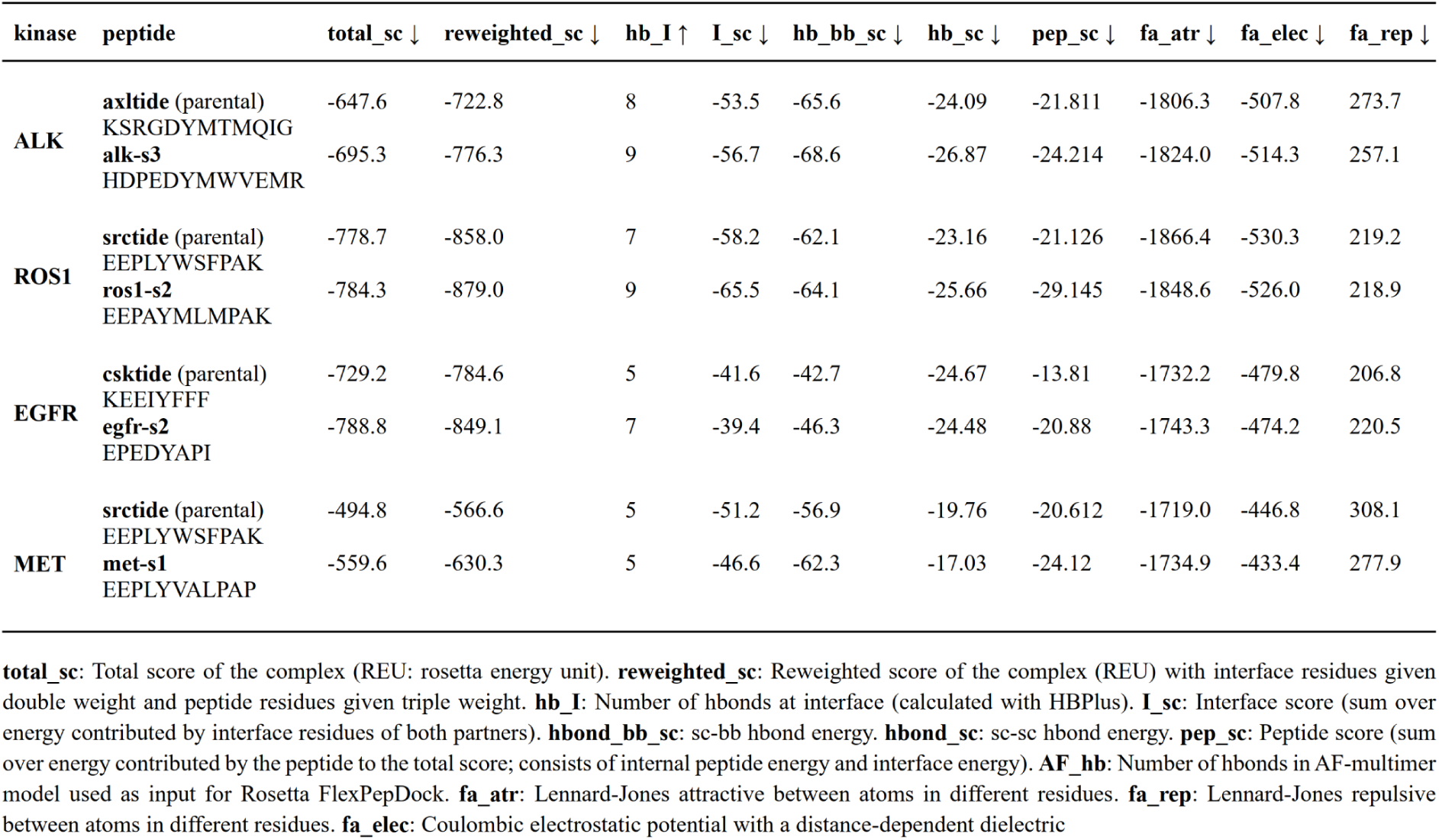
Rosetta FlexPepDock scoring analysis of kinase-peptide complexes.

Figure 5A-D presents structural representations and schematic diagrams highlighting key polar interactions between the kinases and their parental or optimized peptide substrates for ALK, ROS1, MET, and EGFR L858R. By comparing the hydrogen bonding patterns and overall predicted binding poses of the parental peptides with their optimized counterparts, we evaluated potential structural determinants that contribute to the enhanced functional properties. Although the parental peptide MS-axltide (green) forms several hydrogen bonds with residues of ALK (gray) in the structure model shown in Figure 5A, the modeled complex structure with the optimized peptide alk-s3 (yellow) forms slightly different hydrogen bonding patterns, including one additional hydrogen bond with ALK. While many of the hydrogen bonds are retained in the alk-s3, new interactions were made at the substituted peptide residues K1H, G4E, and G12R with kinase residues N151, M199, and N232. These structural changes, particularly the modifications in the hydrogen bonding pattern are predicted to lead to more favorable binding interaction, consistent with the enhanced activity and lower K_m_ observed for ALK-optimized peptides (Figures 2B and 3E).

In the case of ROS1 (Figure 5B), the two additional hydrogen bonds observed in the predicted model of ROS1 kinase with the optimized ros1-s2 substrate were formed by unchanged residues E2, and K11. This indicates that the alterations in the sequences caused some changes in the binding mode of the peptide in a way that optimized the interactions of ROS1 with ros1-s2 for improved binding (Figure 3C) and higher activity (Figure 2D). While the number of hydrogen bonds of the MET complexes of both the parental MS-srctide (green) and optimized met-s1 (yellow) peptide substrates are the same, the unchanged met-s1 peptide residues E2 and Y5 makes new hydrogen bonds with the kinase residues T222 and D137, respectively (Figure 5C).

Furthermore, ros1-s2 residues P3, M8, and K11 form unique hydrogen bonds with ROS1 whereas no corresponding hydrogen bonds exist in met-s1 with MET. Similarly, met-s1 residue E1 makes a unique hydrogen bond with MET compared to the ROS1-ros1-s2 complex. These unique structural differences of the met-s1 and ros1-s2 complexes suggest that the design process introduced specific interactions tailored to the structural features of the binding pocket of the corresponding kinase. This indicates that these unique interactions are likely key determinants of the enhanced selectivity of met-s1 for MET over ROS1 (Figure 4B) and ros1-s2 for ROS1 over MET (Figure 4C).

For EGFR L858R, the unchanged residues E1 and E3 of the optimized egfr-s1 peptide make two new hydrogen bonds with the kinase residues R99 and K209 consistent with the improvement in activity (Figure 5E). Although one hydrogen bond between the parental MS-csktide1 peptide residue E2 and the kinase residue R99 was lost in egfr-s1, the lost bond was compensated for by the new bond between the egfr-s1 peptide residue E1 and the kinase residue K209. These changes in hydrogen bonding pattern suggest potential selectivity of the optimized egfr-s1 peptide for EGFR L858R as opposed to the promiscuous MS-csktide1, which is phosphorylated by several other kinases, including MET, RET M918T, and ROS1^[16]^.

The Rosetta energy and interface scores calculated for each kinase-peptide complex offer a quantitative assessment of the predicted binding interactions of the parental and the most potent designed peptides (Table 1). For all kinases, the Rosetta scores "total_sc" (overall energy of the complex) and "reweighted_sc" (reweighted energy prioritizing interface and peptide residues) of the designed peptides were lower (better) than those of the parental peptides (Table 1), indicating that the designed peptides formed more energetically favorable and stable kinase-peptide interactions. Similarly, the designed alk-s3, ros1-s2, egfr-s2, and met-s1 peptides all have lower (better) Rosetta "pep_sc" (peptide score – overall peptide energy) than their corresponding parental peptides, indicating the designed peptides have better stability and contribute more favorably to the total energy scores of their complexes.

The number of kinase-peptide interface hydrogen bonds (hb_I) for the designed peptides in complex with ALK, ROS1, and EGFR were higher than those of the parental peptides as shown in Table 1 and Figure 5. The increased numbers of polar interactions might play an important role in the improved activity of the designed alk-s3, ros1-s2, and egfr-s2 peptides. However, for MET, the number of interface hydrogen bonds is the same for both the parental and designed peptides. Although the hydrogen bonding pattern differs in the MET complexes of the parental and designed peptides, the improved activity of met-s1 over the parental MS-srctide might not be attributed to just polar interactions. The improvement might also be due to the overall stability of the met-s1 peptide and its complex with MET compared to the parental peptide as indicated by met-s1 having better Rosetta energy scores, including total_sc, reweighted_sc, pep_sc, fa_atr (LJ attractive energy), and fa_rep (LJ repulsive energy) (Table 1)

## 3 Discussion

In this study, we present Subtimizer, a structure-guided computational workflow that approaches the design of potent and selective kinase peptide substrates as a protein design problem. The workflow integrates AF-Multimer for predicting kinase-peptide complex structures, ProteinMPNN for designing novel peptide sequences within this structural context, and structural metrics (ipTM, pTM, ipAE, and pLDDT) for evaluating the confidence and predicted quality of the designed peptides. Unlike previous studies on kinase-substrate relationships that mainly addressed the phosphorylation prediction task, the Subtimizer workflow combines both structure prediction and sequence design tasks to generate optimal peptide substrates for kinases.

We experimentally validated the Subtimizer pipeline by testing designed peptides for five kinases using a luminescence ADP-Glo kinase assay. For four out of the five kinases tested, we successfully identified designed peptides that demonstrated substantially improved kinase activity compared to their parental substrates, with the magnitude of improvement reaching over 300%. Kinetic characterization revealed that the improved activity of the most potent designed peptides for ALK, ROS1, and MET is primarily due to lower K_m_ values (enhanced binding affinity). The improved binding affinity is consistent with the structure-guided design approach, which aims to optimize kinase-peptide interactions within the substrate binding pocket. We also demonstrated that the Subtimizer pipeline generates novel substrates with improved selectivity for the target kinase by showing that substrates designed for ROS1 and MET, while originating from the same parental peptides, exhibited 4-fold and 11-fold improvement in selectivity for their target kinase, respectively. High substrate affinity and selectivity are paramount for developing sensitive kinase assays suitable for low enzyme concentrations and complex biological samples like cell lysates and tissue extracts, as well as direct measurement in vivo ^[7, 18, 20, 21]^.

The computational confidence and quality evaluation steps of the workflow allowed for rapid prioritization of high-confidence designs for experimental validation, as only two to five designed peptides needed to be tested to identify peptides with improved activity. This significantly reduces the experimental burden compared to traditional screening, highlighting both the predictive power and practical efficiency of Subtimizer. Thus, the structure-guided protein design workflow offers a viable approach toward developing the necessary tools to enable comprehensive studies of the human kinome. The ability to generate high-affinity, selective substrates is crucial not only for fundamental research but also for accelerating drug discovery efforts, particularly for the large number of understudied kinases ^[8, 10, 11, 15, 16]^.

While the Subtimizer pipeline demonstrated high success rates for four (ALK, EGFR L858R, MET, and ROS1) of the five kinases tested, designed peptides tested for SRC showed reduced activity compared to the parental MS-srctide peptide. This indicates that the workflow may not work for all kinases, highlighting potential limitations. However, such limitations can be overcome by introducing further improvements to the Subtimizer workflow. The fields of AI-driven protein design and structure prediction are rapidly evolving, with innovative methods emerging frequently. This, in principle, presents an opportunity to enhance the workflow from two angles: by improving the accuracy of (1) the kinase-substrate structure models, and (2) the sequence design step.

We envision that the Subtimizer workflow would be improved by replacing AF-Multimer with the recently published AlphaFold 3 ^[49]^ (or comparable models like Boltz-1 ^[50]^, Protenix ^[51]^, or Chai-1 ^[52]^), the latest iteration of AF that can predict structures of protein complexes with nucleic acids, small molecules, ions, and modified residues. Second, LigandMPNN, the recently developed and ligand-aware version of ProteinMPNN that excels at designing residues near ligands and cofactors ^[53]^, could be used in the sequence design step. Since kinases rely on metal cofactors and nucleotides for phosphorylation, implementing these ligand-aware modifications could improve the performance of the peptide design workflow. For example, such modifications could facilitate the optimization of not only substrate binding but also catalysis as design objectives, potentially overcoming the limitations of K_m_-V*_max_* tradeoff seen with peptides designed for ROS1 and MET (Figure 3C-D).

Another potential limitation of the workflow is a case where the AF-Multimer structure prediction step fails to generate a complex structure that passes the threshold of the confidence metric. In such a case, it might be helpful to either incorporate an automated MD simulation tool for dynamics refinement ^[54]^, or utilize methods such as AlphaRED ^[55]^ that significantly improved the success rate and accuracy of AF-Multimer on difficult cases by integrating ReplicaDock (a physics-based replica exchange docking algorithm).

In summary, this work demonstrates that AI-driven structure-guided protein design holds tremendous potential for scaling up kinase assay development by enabling rapid design of optimal kinase-specific peptide substrates for a larger set of kinases, including high-priority dark kinome members lacking validated substrates. More broadly, results from this study indicate that applications of recent advancements in AI-driven structure-guided protein engineering could generalize to other enzyme-substrate specificity engineering applications beyond kinases, such as proteases ^[46]^ and phosphatases ^[56, 57]^.

## 4 Methods

### 4.1 Workflow for Computational Design of Kinase Peptide Substrates

#### 4.1.1 Kinase-peptide complex prediction

The 3D structures of kinase-peptide complexes were predicted with AF-Multimer ^[32]^ using the ColabFold^[58]^ implementation. The code for local installation of ColabFold was obtained from the repository https://github.com/sokrypton/ColabFold. Sequences of the kinase catalytic domain obtained from UniProt and the starting peptide substrate were used as input to AF-Multimer. For the proof-of-concept study, 25 kinases were each paired with one or more starting peptide substrates that have previously been experimentally validated in-house, making a total of 45 kinase-peptide pairs. For each pair, five rounds of AF-Multimer predictions were run with different seeds, generating five models per round. The number of AF-Multimer recycles and Amber relax cycles were set to 10 and 3, respectively. The top-ranking model from each round was selected, resulting in a total of 5 predictions. Only predictions with an ipTM > 0.75 were considered high-confidence and used for the downstream design step.

#### 4.1.2 Peptide sequence design

The AF-Multimer models were passed to ProteinMPNN, which was used to design novel sequences on the backbone of the starting peptide while keeping the kinase residues fixed and preserving the amino acid identity of the phosphosite (Ser/Thr/Tyr) in the peptide. For the AKT2-gsk3tide complex where two crystal structures are available (PDB IDs 1O6K, 1O6L), these were also included as separate pairs (a total of 47 pairs) for use as input for ProteinMPNN. For each of the input kinase-peptide models, 480 sequences were designed (a total of 2400 for each of the AF-Multimer models) with a sampling temperature of 0.1 and batch size of 32. The designed sequences were clustered using CD-Hit ^[59]^ at 100% sequence identity to eliminate redundant sequences. The ProteinMPNN code was obtained from the repository https://github.com/dauparas/ProteinMPNN.

#### 4.1.3 Structure prediction and evaluation of designed sequences

The newly designed peptide sequences were paired with their target kinase sequences, and their complex structures were predicted again using AF-Multimer with minimal parameters (2 AF recycles, 4 prediction models, and no Amber relaxation). The top-ranking AF-Multimer model for each pair was used as initial guess for the interface prediction and evaluation using a modified version of AF2 (AF2 with-initial-guess) as described by Bennett et al. (2023) ^[47]^. The code for AF2 with-initial-guess interface prediction was obtained from https://github.com/nrbennet/dl_binder_design. The ipAE, ipTM, pTM, and pLDDT scores were used to evaluate and rank the ProteinMPNN-generated sequences to identify a set of high-confidence designs for experimental validation. For the kinases (ALK, MET, ROS1, EGFR L858R, and SRC) selected for experimental validation, two to five designed peptides with low ipAE and high ipTM and pLDDT scores were chosen for experimental tests.

#### 4.1.4 Structural analysis and energy evaluation

Structure refinement and quantitative assessment of predicted binding interactions was performed using the Rosetta FlexPepDock protocol ^[48]^. AF-Multimer models of the kinase complexes of the most potent designed peptide and their parental counterparts were used as input for FlexPepDock refinement and scoring using the Rosetta energy function. The refined structure with the lowest total Rosetta energy score was used for interaction analysis. Hydrogen bond interactions between kinase and peptide residues were calculated with HBPLUS 3.2 ^[60]^ using angle and distance criteria of D-H-A > 125°, D-A < 3.45 Å. Predicted kinase-peptide complex structures and key interactions were visualized using PyMOL (Schrödinger, LLC). Plots were generated using Python or GraphPad Prism.

### 4.2 ADP-Glo Kinase Assays

ADP-Glo Kit (ADP-Glo™ Kinase Assay, Cat#V6930) was obtained from Promega. The kinases ALK (#08-518), MET (#08-151), ROS1 (#08-163), and EGFR L858R (#08-502) were products of Carna Biosciences. SRC was expressed and purified as previously reported ^[61, 62]^. Substrate peptides were obtained from Biomatik Corporation (Wilmington, DE, USA). All components were equilibrated to 25°C prior to setting up reactions in 384-well microplates (White ProxiPlate 384-shallow well Plus). Final working concentrations were enzyme: 1.25-10 nM, ATP 10 *µ*M, peptide 1 *µ*M. All reagents were added manually and incubated for 90 minutes. Experiments were repeated twice with results expressed as mean ± standard deviation. Luminescence was detected on a Synergy Neo2 plate reader.

### 4.3 LIMS-Kinase Assays – Enzyme Optimization

All components were equilibrated to 25°C prior to setting up reactions in 384-well microplates (Costar 3657 round bottom polypropylene). All reagents were added manually, and final working concentrations were enzyme 0.63-10 nM, ATP 100 *µ*M, peptide 1-10 *µ*M. Enzyme was first added to plate, followed by peptide/ATP mix in buffer to initiate reaction. Reactions were quenched at 20, 40, 60, 80, 100 minutes with formic acid at final concentration of 1%. Experiments were repeated twice and results expressed as mean ± standard deviation.

### 4.4 LIMS-Kinase Assays – Kinetic Analysis

Final working concentrations of enzyme varied by kinase. ATP was at 100 *µ*M, and peptide concentrations started at 300-400 *µ*M followed by 2-fold or 3-fold dilutions. All components were equilibrated to 25°C prior to setting up reactions. Peptide dilutions were added to 384-well microplates (Costar 3657 round bottom polypropylene). All reagents were added manually. Buffer, enzyme, and ATP mixture were added to initiate reaction. Reactions were quenched at set time-points with 1% formic acid (final). Activity was evaluated by measuring MS signal of ADP-specific fragment ion. Experiments were repeated twice, with results expressed as mean ± standard deviation.

### 4.5 Calculation of **K_m_**

K_m_ was evaluated by linear regression of the first three time-points at each concentration of peptide. The slope was used to represent initial velocity in graph of initial velocity versus concentration of peptide. Nonlinear regression using GraphPad Prism’s Michaelis Menten equation was used to estimate the K_m_.

### 4.6 Peptide Selectivity Assays

For MET selectivity assay, reagent final working concentrations were MET 5 nM, peptides 134 *µ*M (K_m_ for met-s1 with MET), and ATP 100 *µ*M. For ROS1 selectivity assay, reagent final working concentrations were ROS1 0.63 nM, peptides 20.3 *µ*M (K_m_ for ros1-s2 with ROS1), and ATP 100 *µ*M. All components equilibrated to 25°C prior to setting up reactions. All reagents added manually. Peptides added to 384-well microplates (Costar 3657 round bottom polypropylene). A pre-mixed buffer, enzyme, and ATP solution was added to initiate reaction. Reaction was quenched with 1% formic acid (final) following 30-minute incubation at room temperature. Kinase activity was evaluated by measuring MS signal of ADP-specific fragment ion. Experiments were repeated twice, with results expressed as mean ± standard deviation. Time 0 (background) values were subtracted from subsequent time-points.

### 4.7 RapidFire Chromatography and Mass Spectrometry

RapidFire liquid chromatography and mass spectrometry methods were performed as previously described^[16]^.

### 4.8 Statistical Analysis

All experiments were performed in duplicate with results expressed as mean ± standard deviation. Time 0 background values were subtracted from subsequent time-points where indicated.

## 5 Code Availability

The code of the Subtimizer pipeline is publicly available at https://github.com/abeebyekeen/subtimizer.

## Acknowledgements

We thank the University of Texas Southwestern Medical Center and the Simmons Comprehensive Cancer Center for institutional support. This work was supported by Welch Foundation I-1829 (K.D.W), NIH UM1CA294119 (K.D.W), P50CA070907 (K.D.W) and P30CA142543.

---

**Figure S1:**
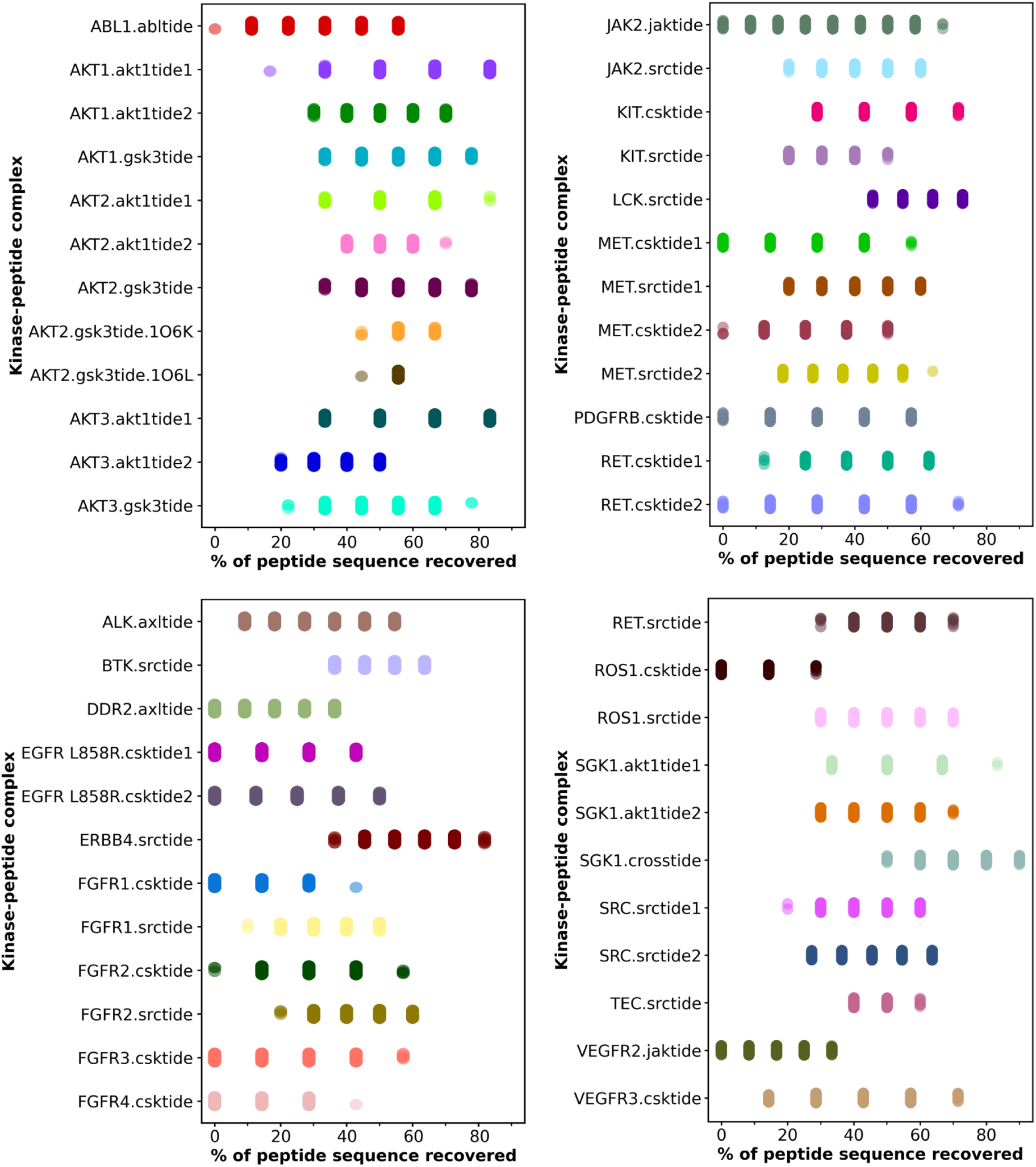
Substrate sequence recovery analysis across kinase families. Percentage of peptide sequence recovery for all 46 kinase-substrate pairs tested in the validation study. Each dot represents recovery percentage for individual Subtimizer designs. Most kinase-substrate pairs (>50%) achieve *≥*70% sequence recovery, demonstrating robust identification of functionally important residues.

**Figure S2:**
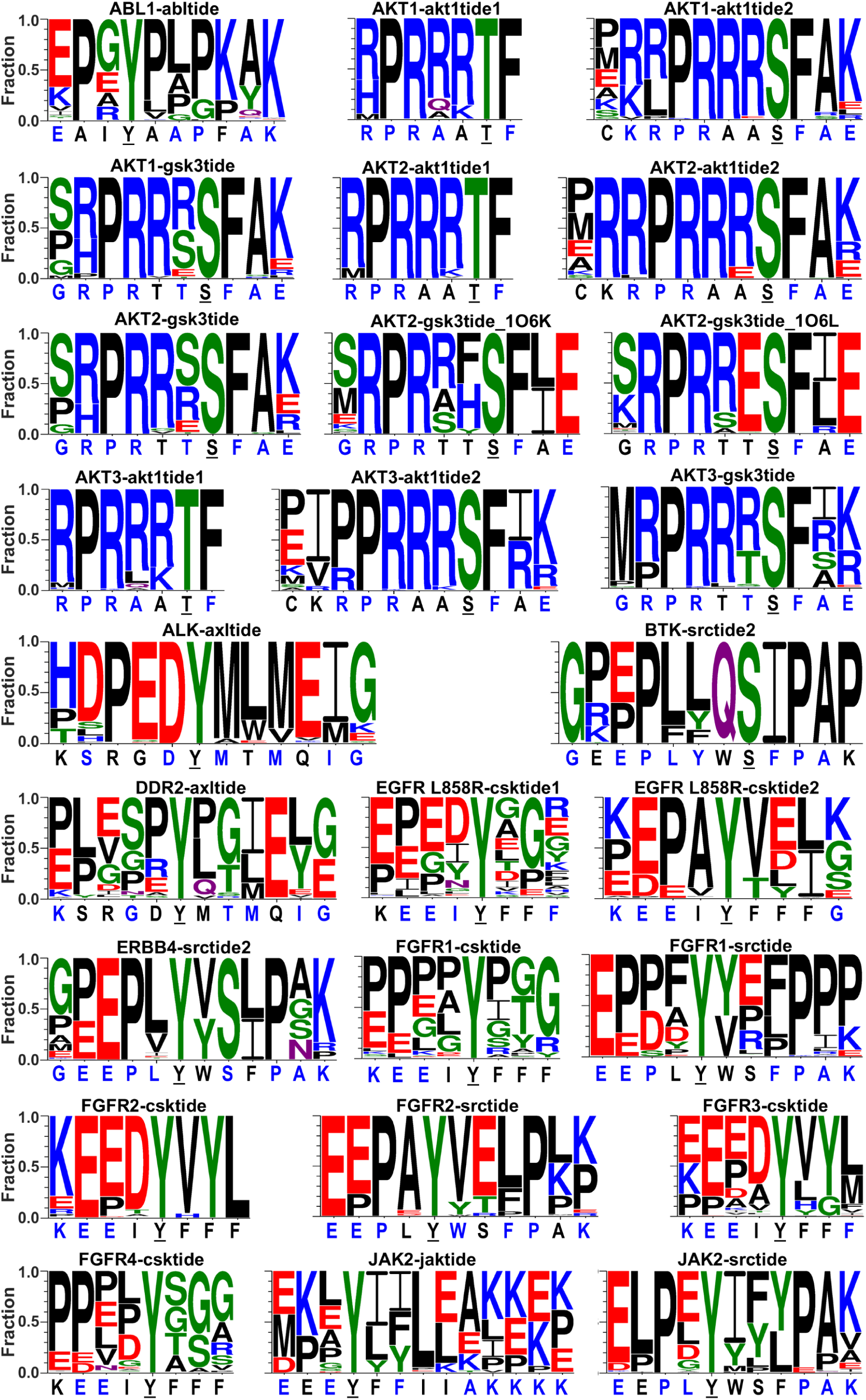
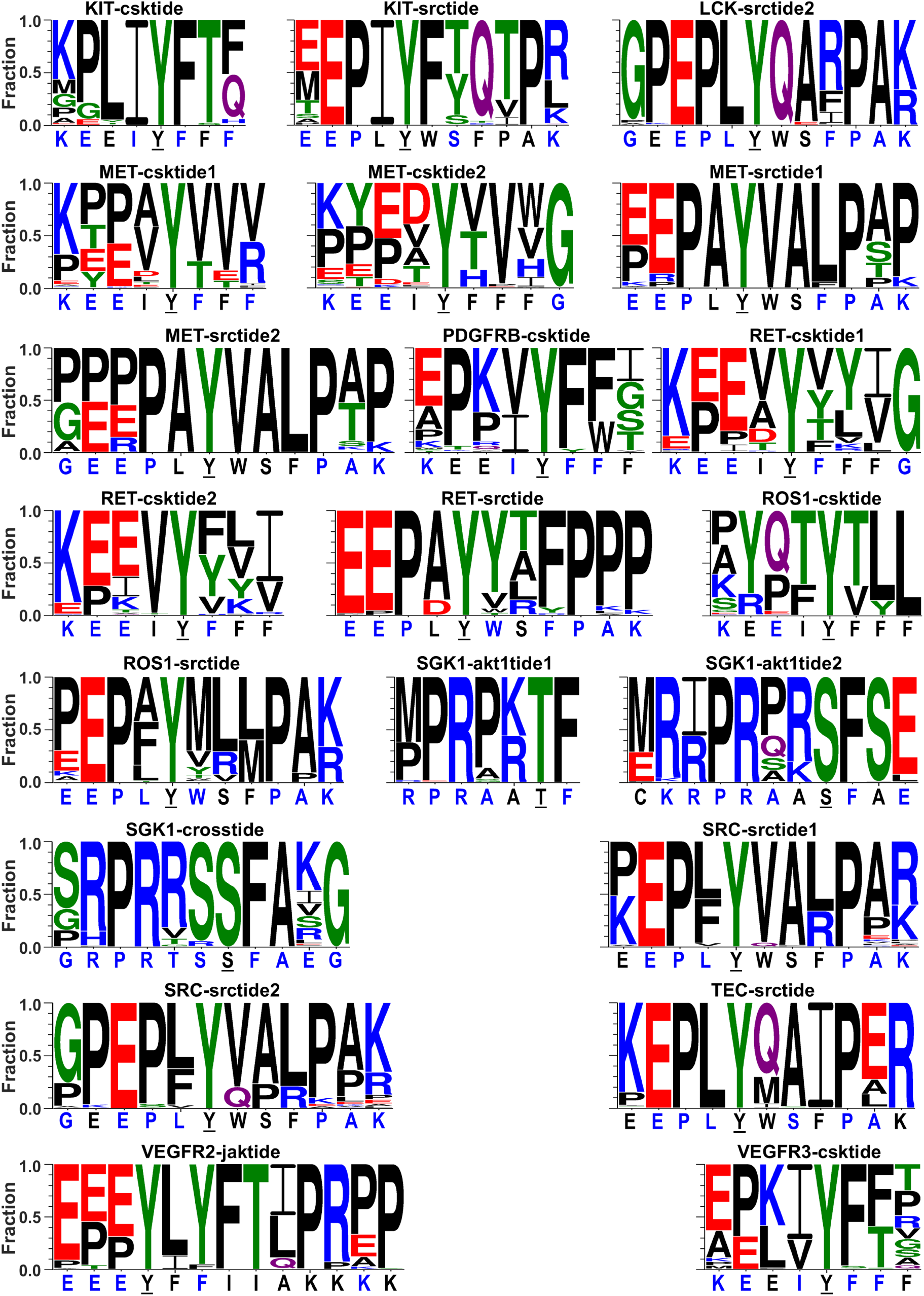
Sequence logos of Subtimizer-designed peptides. Amino acid frequency and preferences at each position for designed peptides across kinase-substrate pairs. Letter height corresponds to amino acid frequency. Phosphorylatable residues and structurally important positions show strong conservation, while variable positions indicate regions of sequence optimization.

**Figure S3:**
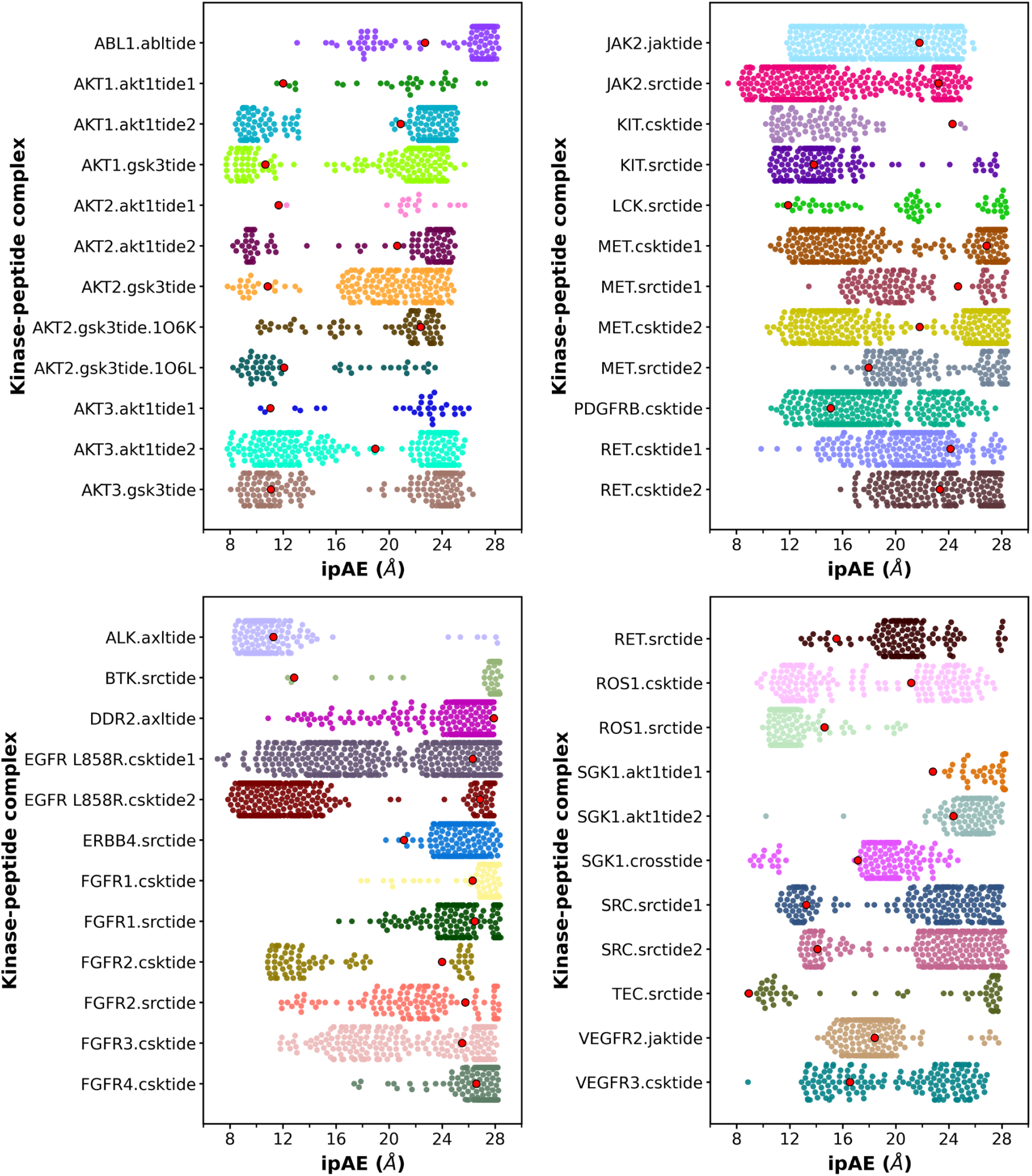
Distribution of interface predicted aligned error (ipAE) scores for all 47 kinase-substrate pairs. The ipAE scores of Subtimizer-designed peptides were compared with parental peptides (red dots). For 46 of the 47 pairs (98%), Subtimizer generated peptides with lower ipAE scores than parental substrates, with most achieving scores below 10 Å.

**Figure S4:**
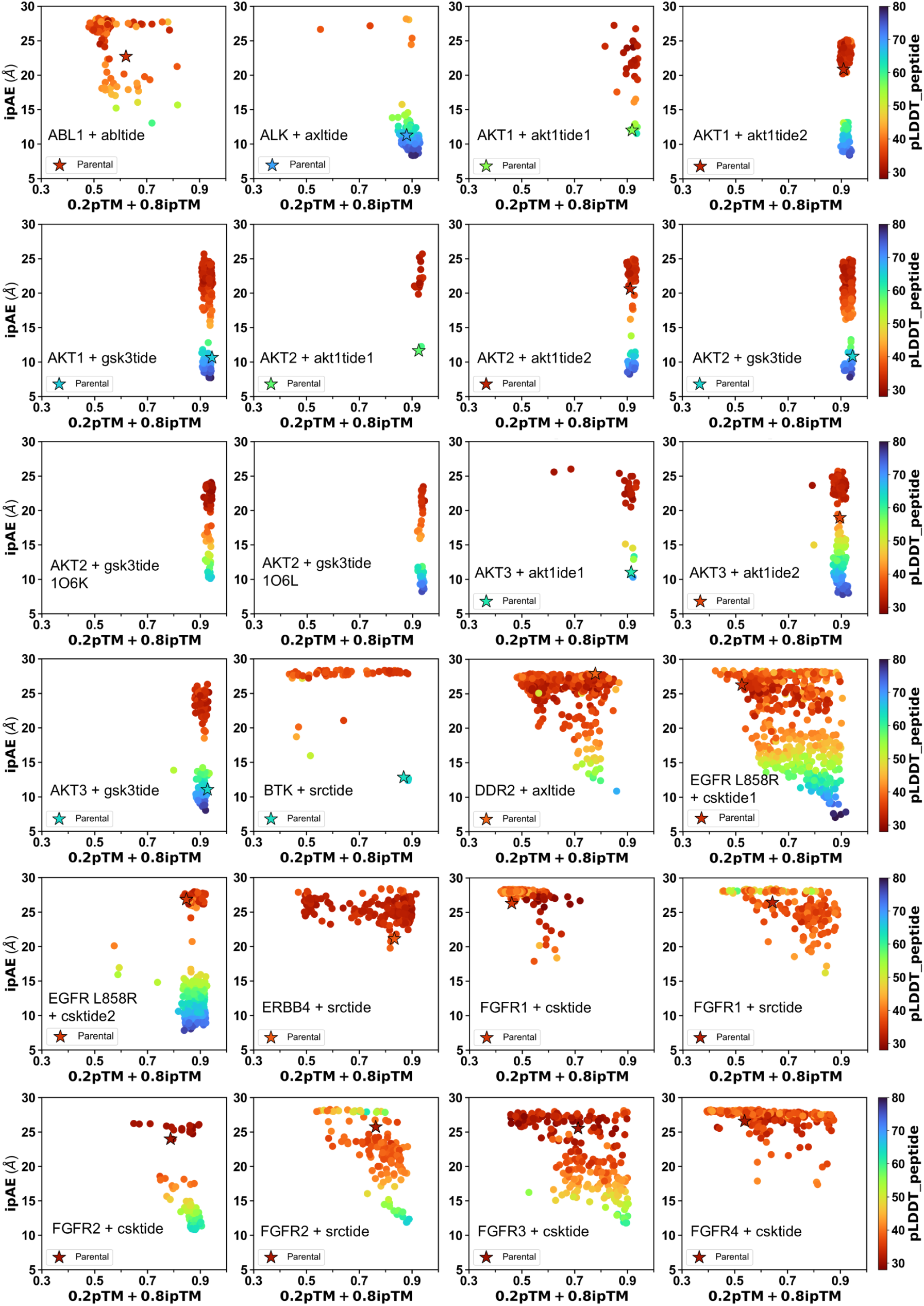
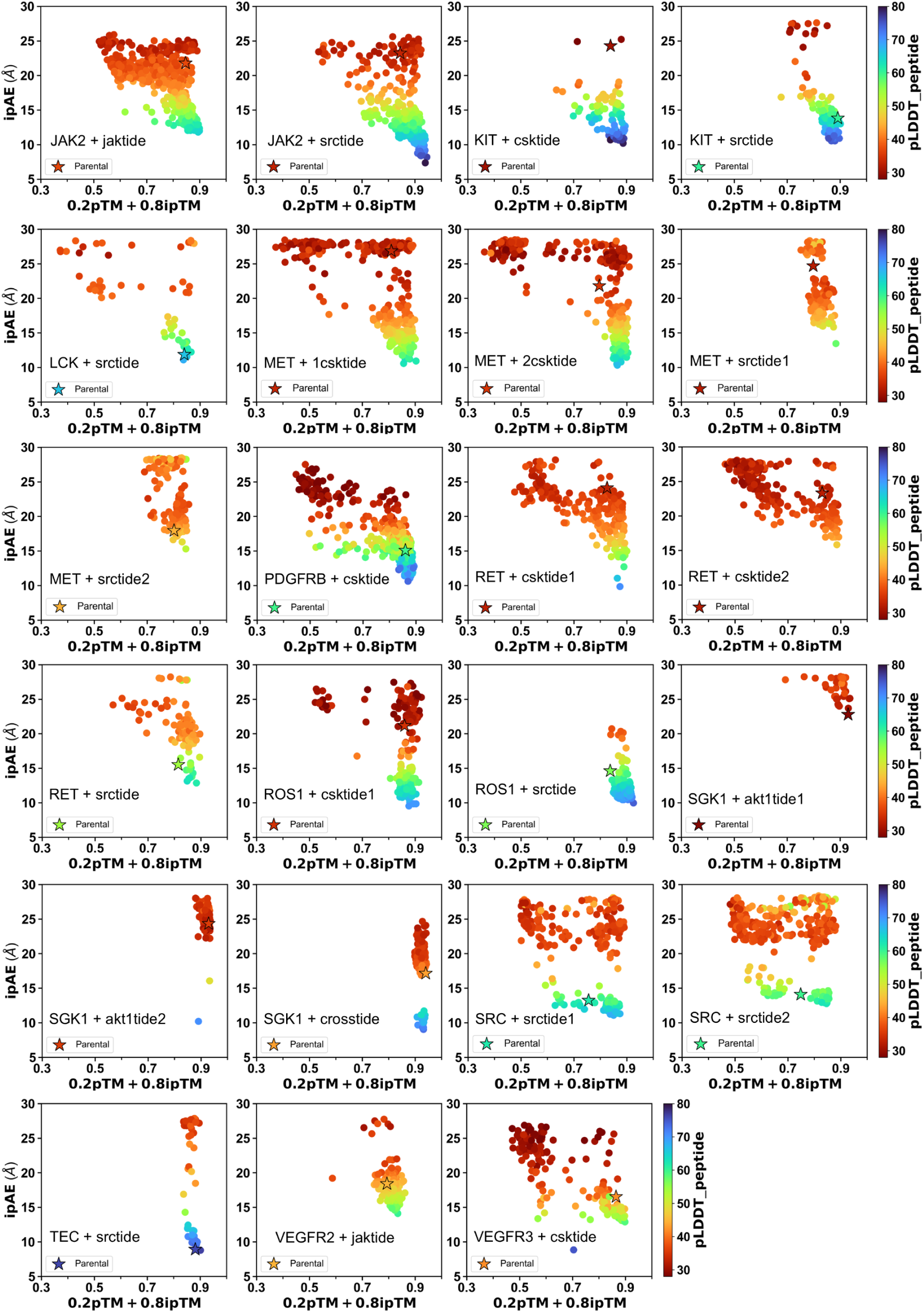
Correlation between structural evaluation metrics. Scatter plots showing relationship between ipAE and combined confidence scores (0.2 *·* pTM + 0.8 *·* ipTM) for all kinase-substrate pairs. Color gradient of dots show peptide confidence (pLDDT). Strong inverse correlation validates the use of these metrics for design evaluation.

**Figure S5:**
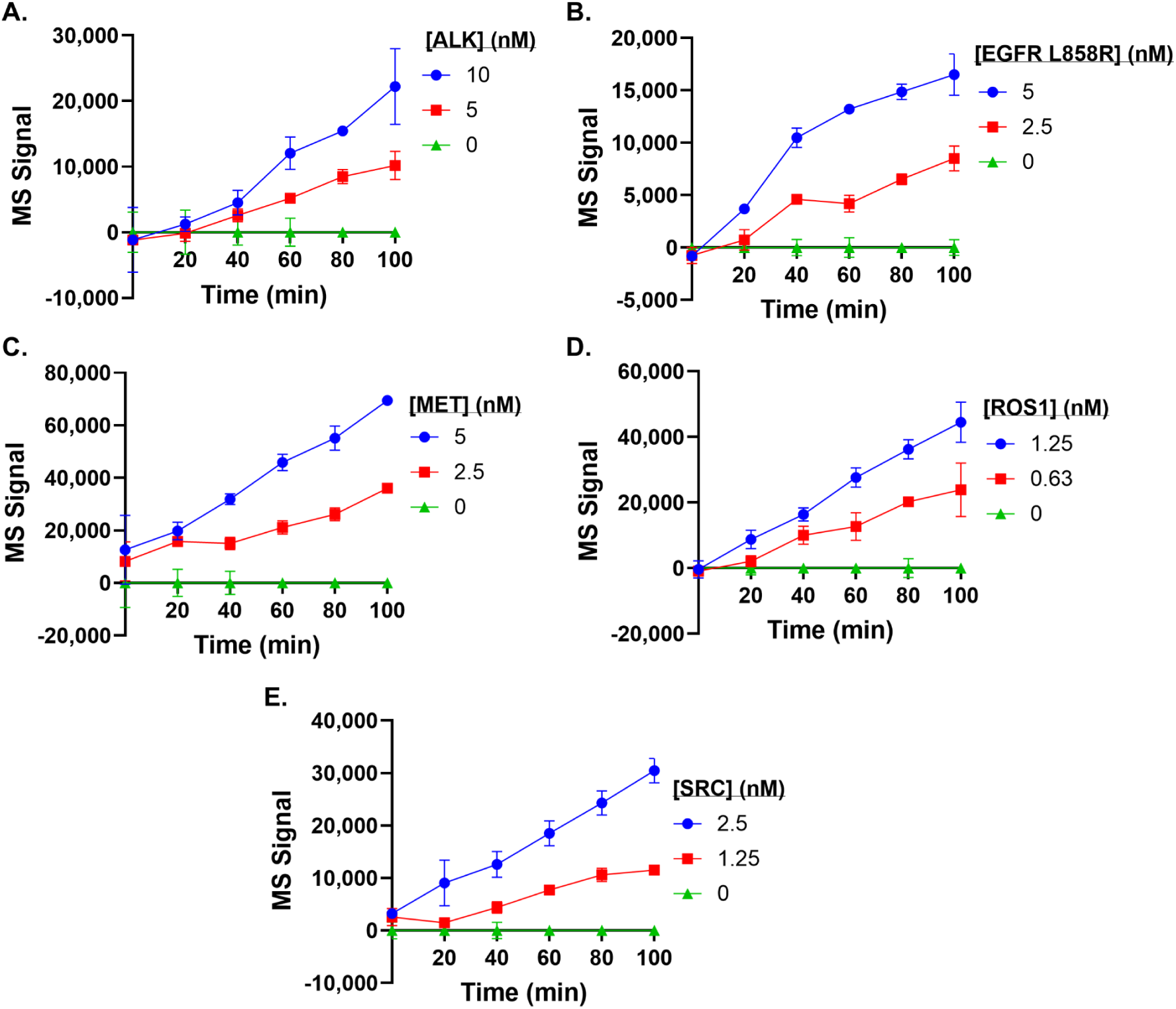
LIMS-kinase assay optimization confirms enzyme concentration- and time-dependent ADP production. Time-course analysis of ADP production across different enzyme concentrations for all five kinases tested, validating functional assay conditions for kinetic analysis. All assays performed with 100 *µ*M ATP at varying peptide concentrations. Green triangles represent background controls (0 nM enzyme) showing minimal non-enzymatic ADP production. (**A**) ALK with 10 *µ*M peptide substrate. (**B**) EGFR L858R with 1 *µ*M substrate. (**C**) MET with 5 *µ*M substrate. (**D**) ROS1 with 10 *µ*M substrate. (**E**) SRC with 5 *µ*M substrate.

**Figure S6:**
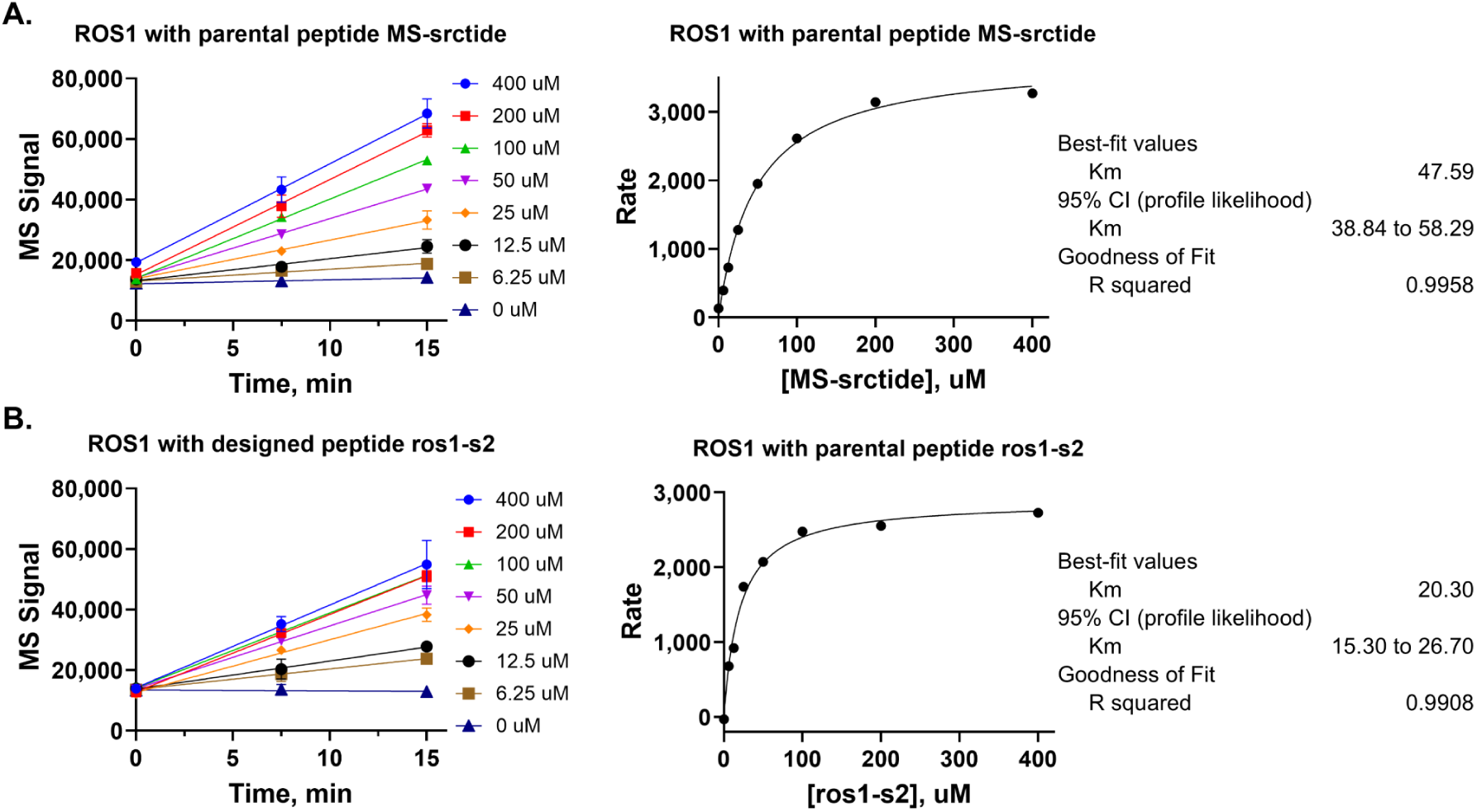
Detailed kinetic analysis of ROS1. (**A**) ROS1 kinetic analysis with parental peptide MS-srctide showing time-course measurements and Michaelis-Menten analysis (K_m_ = 47.59 *µ*M, R^2^ = 0.9958). (**B**) ROS1 analysis with designed peptide ros1-s2 (K_m_ = 20.30 *µ*M, R^2^ = 0.9908), demonstrating >2-fold improvement in apparent binding affinity.

**Figure S7:**
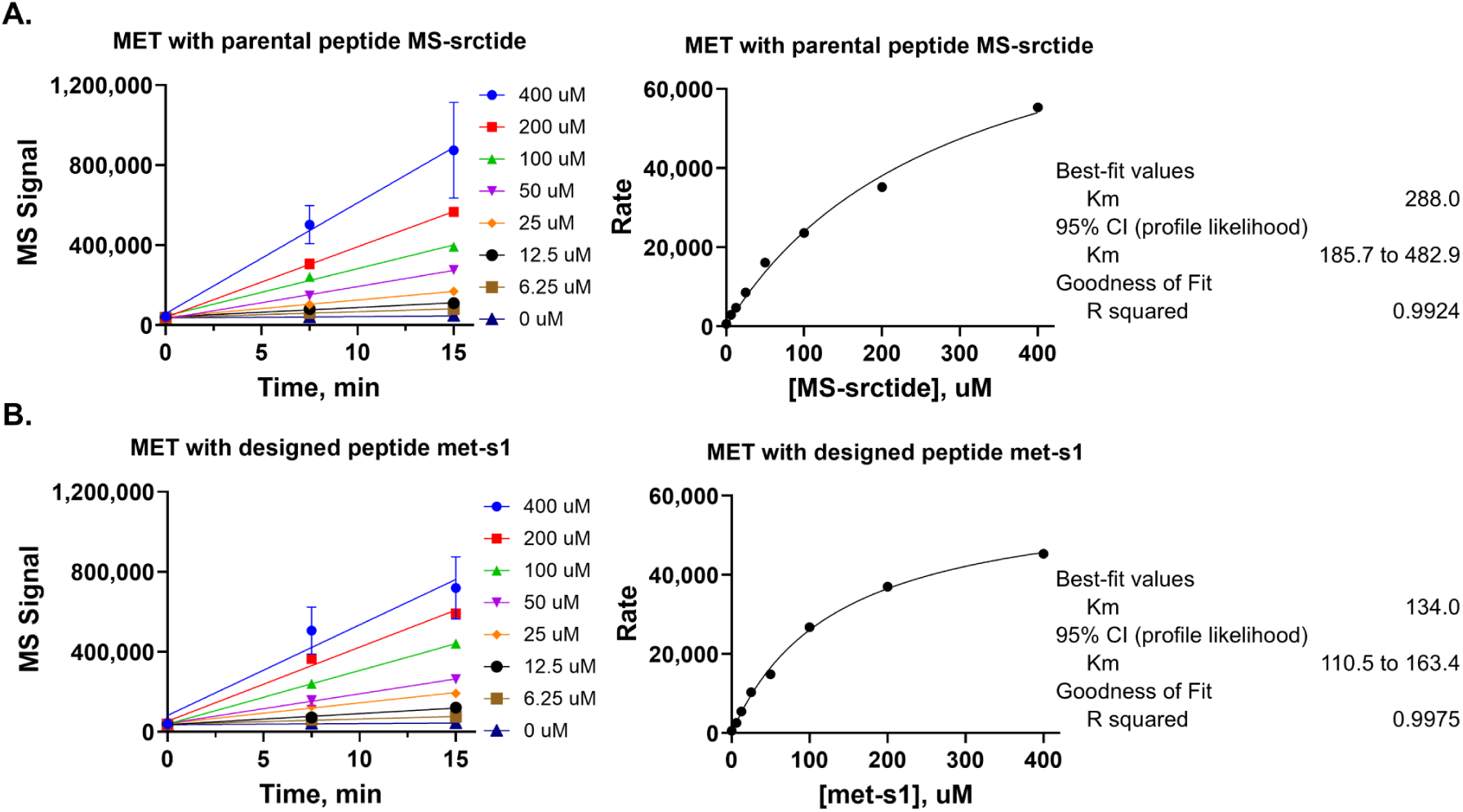
Kinetic analysis of MET kinase. (**A**) MET kinetic analysis with parental peptide MS-srctide showing concentration-dependent kinetics. (**B**) MET analysis with designed peptide met-s1, revealing improved saturation kinetics with >2-fold reduction in K_m_ (288 *µ*M to 134 *µ*M).

**Figure S8:**
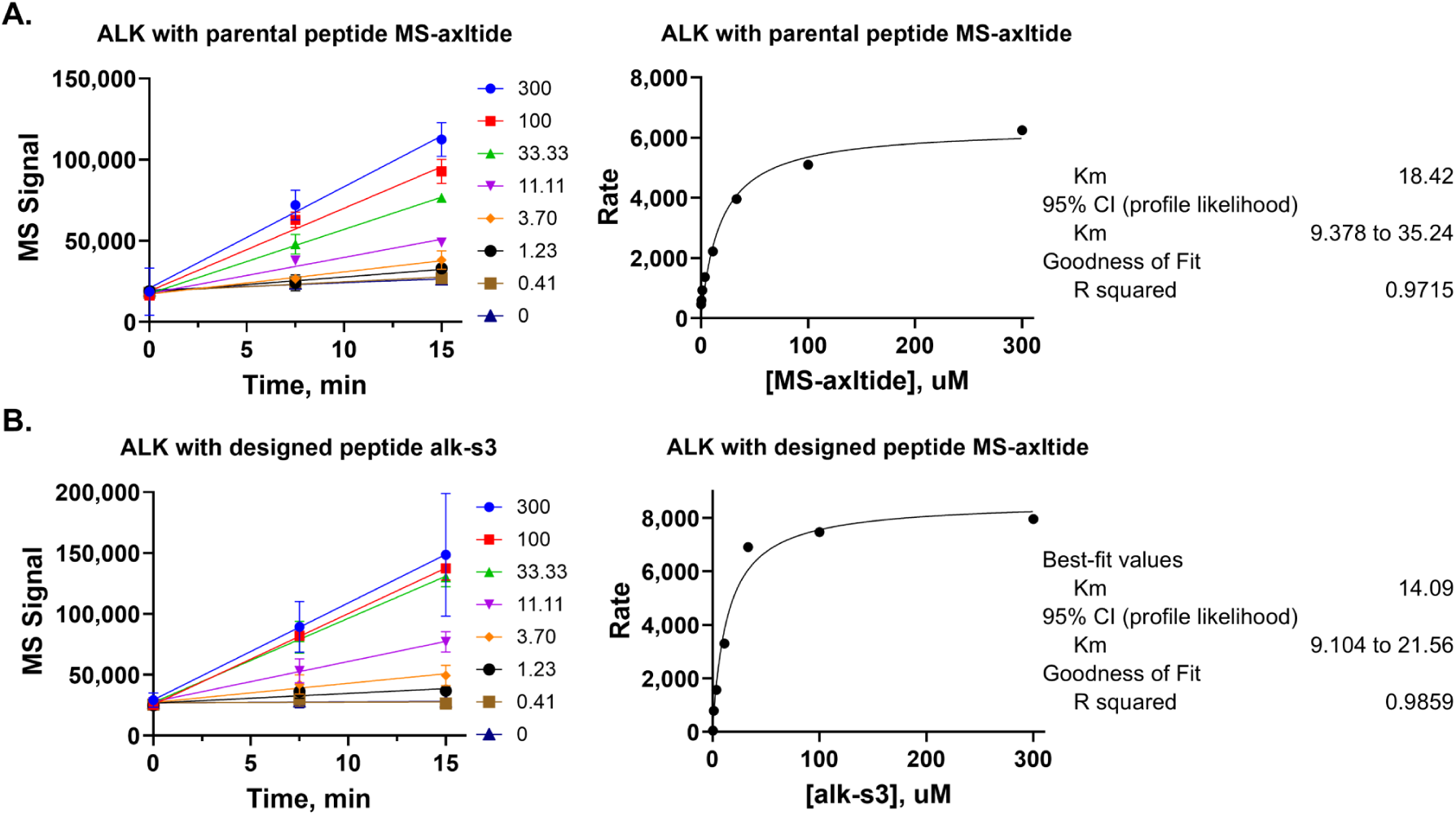
Kinetic characterization of ALK kinase. (**A**) ALK kinetic analysis with parental peptide MS-axltide. (**B**) ALK analysis with designed peptide alk-s3 showing improved binding affinity (K_m_: 18.4 *µ*M to 14.1 *µ*M) and enhanced catalytic efficiency.

**Figure S9:**
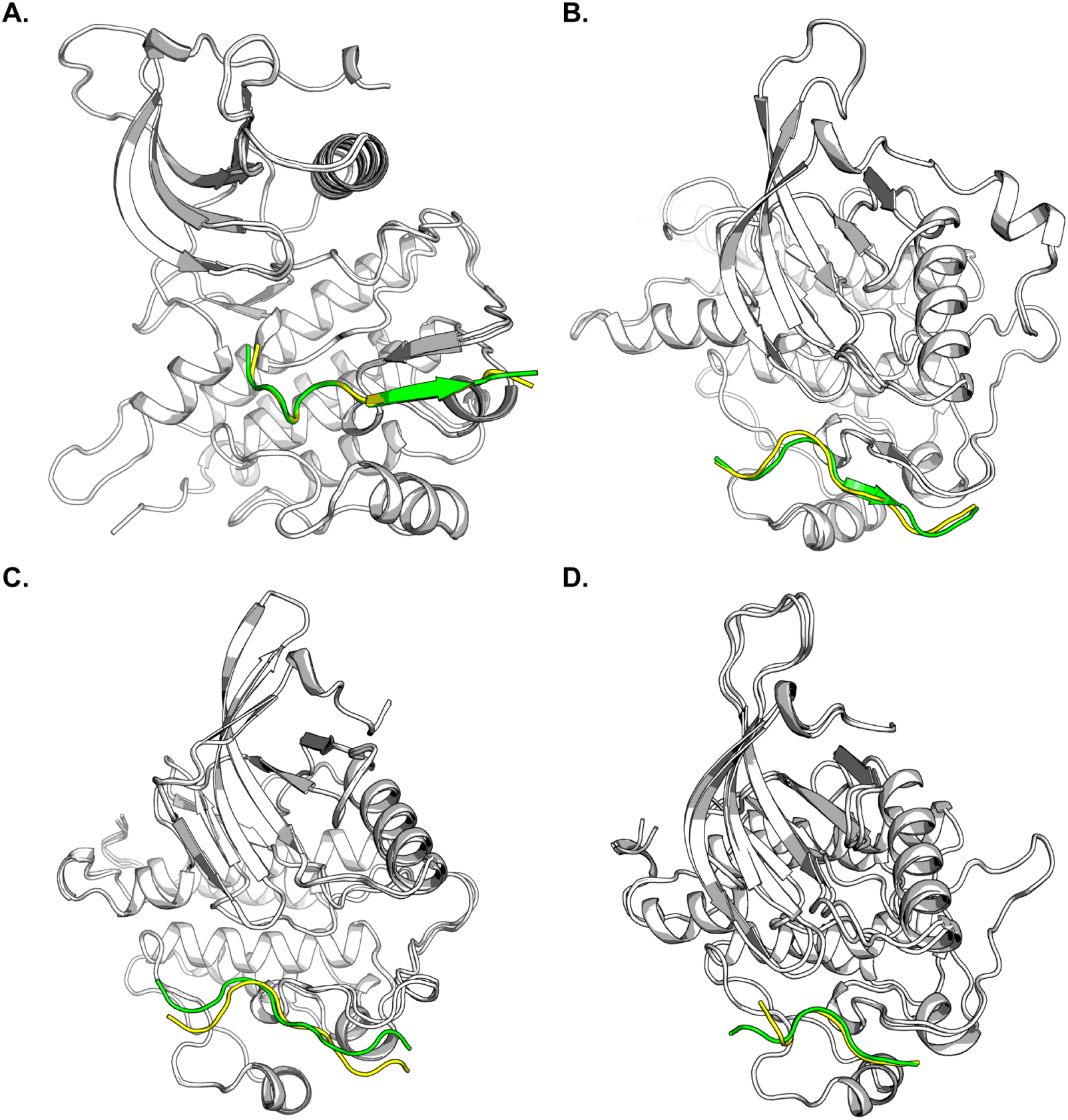
Structural comparison of AlphaFold-Multimer predicted structures of kinase-peptide complexes showing parental substrates (green) and Subtimizer-designed peptides (yellow) bound to the kinases (gray) ALK (**A**), ROS1 (**B**), MET (**C**), and EGFR L858R (**D**). Overlays reveal conformational differences and optimized binding modes in designed peptides.

## Notes

### Competing Interest Statement

K.D.W. has received consulting fees from Sanofi Oncology, Amgen, AstraZeneca, Boehringer Ingelheim and is a member of the SAB for Vibliome Therapeutics and Vellorum Pharmaceuticals. K.D.W. has received research funding from Revolution Medicines and Elekta. K.D.W. is a co-founder and equity holder in Stabilix LLC. K.D.W. declares that none of these relationships are directly or indirectly related to the content of this manuscript.

